# SVclone: inferring structural variant cancer cell fraction

**DOI:** 10.1101/172486

**Authors:** Marek Cmero, Cheng Soon Ong, Ke Yuan, Jan Schröder, Kangbo Mo, PCAWG Evolution and Heterogeneity Working Group, Niall M. Corcoran, Tony Papenfuss, Christopher M. Hovens, Florian Markowetz, Geoff Macintyre

**Author notes:** Current address: School of Computing Science, University of Glasgow, Sir Alwyn Williams Building, Glasgow, G12 8RZ, UK. Members of the PCAWG Evolution and Heterogeneity Working Group: Pavana Anur, Rameen Beroukhim, Paul Boutros, David D. Bowtell, Peter Campbell, Elizabeth L. Christie, Marek Cmero, Yupeng Cun, Kevin Dawson, Jonas Demeulemeester, Stefan C. Dentro, Amit Deshwar, Nilgun Donmez, Roland Eils, Yu Fan, Matthew Fittall, Dale W. Garsed, Moritz Gerstung, Gad Getz, Santiago Gonzalez, Gavin Ha, Kerstin Haase, Marcin Imielinski, Yuan Ji, Clemency Jolly, Kortine A. Kleinheinz, Juhee Lee, Henry Lee-Six, Ignaty Leshchiner, Dimitri G. Livitz, Geoff Macintyre, Salem Malikic, Florian Markowetz, Inigo Martincorena, Thomas J. Mitchell, Quaid Morris, Ville Mustonen, Layla Oesper, Martin Peifer, Myron Peto, Benjamin J. Raphael, Daniel Rosebrock, Yulia Rubanova, S. Cenk Sahinalp, Adriana Salcedo, Matthias Schlesner, Steve Schumacher, Subhajit Sengupta, Paul T. Spellman, Lincoln Stein, Maxime Tarabichi, Peter Van Loo, Ignacio Vázquez-García, Shankar Vembu, Wenyi Wang, David C. Wedge, David A. Wheeler, Jeff Wintersinger, Tsun-Po Yang, Xiaotong Yao, Fouad Yousif, Kaixian Yu, Ke Yuan, Hongtu Zhu. Correspondence to: Geoff Macintyre and Marek Cmero.

## Abstract

We present SVclone, a computational method for inferring the cancer cell fraction of structural variant breakpoints from whole-genome sequencing data. We validate our approach using simulated and real tumour samples, and demonstrate its utility on 2,658 whole-genome sequenced tumours. We find a subset of liver, breast and ovarian cancer cases with decreased overall survival that have subclonally enriched copy-number neutral rearrangements, an observation that could not be discovered with currently available methods.

**One Sentence Summary:** SVclone is a novel computational method for inferring the cancer cell fraction of tumour structural variation from whole-genome sequencing data.

---

The clonal theory of cancer evolution^1^ posits that cancers arise from a single progenitor cell that has acquired mutations conferring selective advantage, resulting in the expansion of a genetically identical cell population or “clone”. As this cancer grows, a process akin to Darwinian species evolution emerges with subsequent genetically distinct populations arising from the founding clone via the continual acquisition of advantageous genomic aberrations. Consequently, tumours are likely to consist of a genetically heterogeneous combination of multiple cell populations, the extent of which has been revealed through the use of whole-genome sequencing^2,3^. As clones can respond differently to therapy^4^, understanding this cellular diversity has important clinical implications^5^.

The mutations belonging to each clone in a tumour can be interrogated using bulk whole-genome sequencing, with mutation detection subject to sequencing depth, tumour cellularity, clonality and mutation copy-number^6^. The expansion of each clone over the life of a tumour is encoded in the allele frequency of somatic mutations^7^. In order to model clonal expansions, the variant allele frequency (VAF) must be converted to a cancer cell fraction (CCF), the fraction of cancer cells within which the variant is present. Events appearing inr all cancer cells (CCF 100%) are considered clonal and due to a pervasive expansion. Events appearing in a subset of cells (CCF <100%) are considered subclonal and part of an ongoing expansion. Estimating the cancer cell fraction of events is challenging, as the observed variant allele frequency depends on the amount of normal cell admixture (purity) and local copy-number.

Given these challenges, previous computational approaches for estimating CCF have focused on individual facets of this complexity, commonly limiting their view to single nucleotide variants (SNVs)^8-13^ or somatic copy-number aberrations (SCNAs)^14-16^. This has left the clonality of balanced rearrangements largely unexplored, despite their implication in oncogenic fusions^17^ and subclonal translocations conferring a drug resistant phenotype^18^. While SNV-based approaches have provided solutions to the problem of downstream inference of mutation CCF, they cannot be used for structural variant (SV) breakpoint data as: (i) no complete and robust methodology exists yet to calculate VAFs from SVs (Fan et al.^19^ provides a limited framework that does not correct for DNA-gains or support all SV types); (ii) the handling of copy-number changes for SNVs does not translate cleanly to SV data (additionally, SVs can themselves cause copy-number changes); and (iii) due to the relatively small number of data points (on average compared to SNVs), false positive SVs greatly diminish clustering performance, hence a robust filtering methodology is required to consider only high-confidence SVs.

To address this gap, we have developed SVclone, an algorithmic approach that infers and clusters CCFs of SV breakpoints. It considers all types of large-scale structural variation (SV), including copy-number aberrant and copy-number neutral variation. The SVclone pipeline and a summary of its performance validation is shown in Figure 1a. Detailed explanations for each step can be found in the Methods and Supplementary Information. The SVclone algorithm consists of five steps: annotate, count, filter, cluster and post-assign. The annotate, count and filter steps are used to obtain supporting and normal (non-supporting) reads of high-confidence SVs for clustering. The clustering step jointly infers CCF and groups variants of similar CCF. The post-assign step assigns the remaining variants a most-likely CCF, given the clusters obtained from the previous step. SV calls are required as input into the annotate step (single-nucleotide resolution paired SV loci), and the corresponding whole-genome sequencing file in BAM format.

**Fig. 1.**
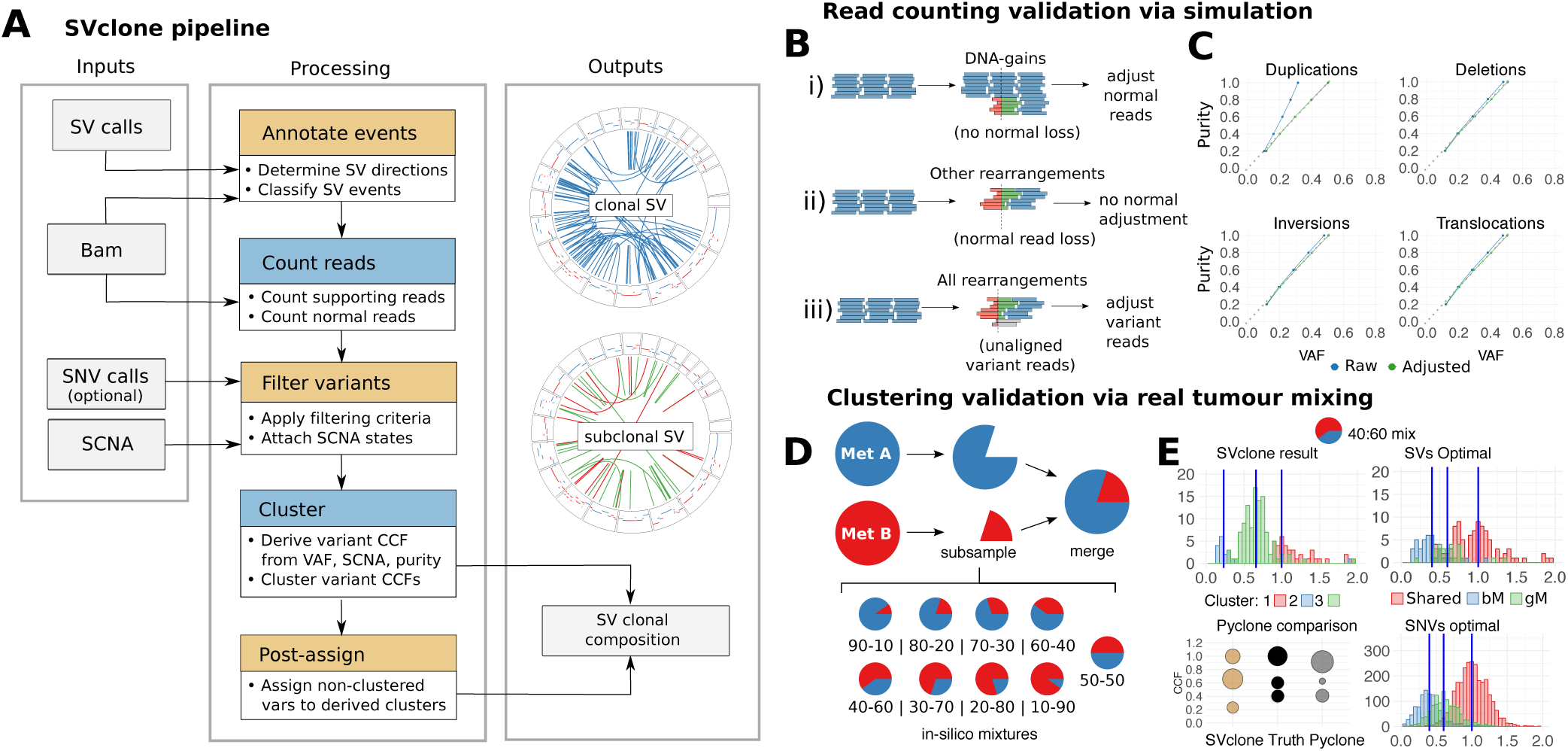
**A)** Flow-chart of the SVclone pipeline. **B)** Schematic showing adjustments required for different classes of rearrangements. **C)** Effect of adjusting raw VAFs at purity levels at 20% to 100% in 20% increments, where the expected VAF is half the purity level (black line). **D)** Schematic illustrating subsampling and merging process to create in-silico mixtures of real tumour samples. **E)** Example results from the 40:60 mixture (blue lines are the cluster mean CCFs). PyClone comparison shows circle size representing the proportion of variants belonging to cluster, while circle centre represents cluster CCF.

The structural variant allele frequency calculation in SVclone depends greatly on high quality read counts for normal and variant DNA. Therefore, to test if any potential bias could affect these counts, we simulated reads from SVs with known allele frequency and computed the observed VAF. Figure 1c shows that for each of the SV classes the observed VAF is systematically underestimated. The lower variant read count causing this discrepancy is due to a small number of variant reads not being aligned (Figure 1b). Duplications are more pronounced as they also have an increased normal read count due to DNA gains showing no loss of normal DNA (Figure 1b). To account for this bias, SVclone employs a simple scaling factor that incorporates tumour purity to calibrate the supporting read counts and infer the true underlying VAF (Figures 1b and 1c).

To test the ability of SVclone to infer clonal expansion frequencies using SVs we created a set of samples with known clonal frequencies by subsampling and mixing read data in 10% increments from two previously sequenced prostate cancer metastases from a single patient^20^ (Figure 1d). The prostate cancer samples used to create the mixtures had no evidence of subclonality (Hong et al.^20^ Supplementary Figure 2d), and had similar coverage and tumour purity. SVclone applied to these samples was able to identify the correct number of clusters in all cases (Supplementary Figure 4). The average mean squared error for cluster frequencies was 0.0109. The average adjusted Rand index was 0.1445, generally low, indicating that while SVclone can correctly identify the number of clusters and closely estimate their respective CCFs, correct assignment of SVs to their true clusters remains a challenge. To provide some insight into this we calculated optimal CCF distributions for SVs (a transformation of the VAF given the most likely copy-number state of each SV if the ‘correct’ cluster mean is known). The overlapping and multi-modal distributions in Figure 1e clearly illustrate the difficulty of the problem when considering breakpoint data, especially compared to optimal distributions for SNVs (Figure 1e) which are more separable.

We also compared SVclone’s performance (run on SVs) to PyClone^9^, a state of the art clustering approach (run on SNVs). SVclone and PyClone results were in good concordance, with PyClone finding 3 clusters in 5/9 mixtures and SVclone in 9/9 mixtures (Figure 1e and Supplementary Figure 4). SVclone’s mean squared error (MSE) across all mixture proportions was 0.011, compared with with PyClone’s MSE of 0.003. A higher deviation from the true cluster mean is expected, given SVclone has comparatively fewer variants available for clustering. SVclone’s comparable performance indicates that clonal structure can be effectively reconstructed with high concordance and accuracy, despite the relative deficit in variant number. Both SVclone and PyClone had little difficulty in identifying minor subclonal clusters in the mixtures due to the lack of overlapping CCFs from clonal or major subclonal clusters. One major advantage of using SVclone is that it can also be used in coclustering mode, where CCF estimates can be given for SVs and SNVs simultaneously, thus substantially increasing the number of variants available for clustering.

We applied SVclone to 2,658 WGS samples from the pan-cancer analysis of whole genomes (PCAWG) project (dcc.icgc.org/pcawg), coclustering both SVs and SNVs, an extensive discussion of which can be found in the PCAWG-11 heterogeneity manuscript (Dentro et al. in preparation). Here, downstream analysis was performed on the 23 tumour types showing ≥ 20 samples with >10 SVs and SNVs (n=1,773). A comparison of the fraction of subclonal SVs versus SNVs showed different patterns across tumour types (Figure 2a). Briefly, tumour types showing more subclonal SVs versus SNVs include 80.4% of lung squamous cell carcinomas and 77.4% of osteosarcomas. This is in contrast to the 36.3% of lung adenocarcinomas and 40% of gastric cancers (Supplementary Table 3). Some cancers contained subsets with distinct patterns of clonality, for instance liver cancers contained a cluster of 21 samples with high SV subclonality (≥ 50%) and low SNV subclonality (< 30%).

**Fig. 2.**
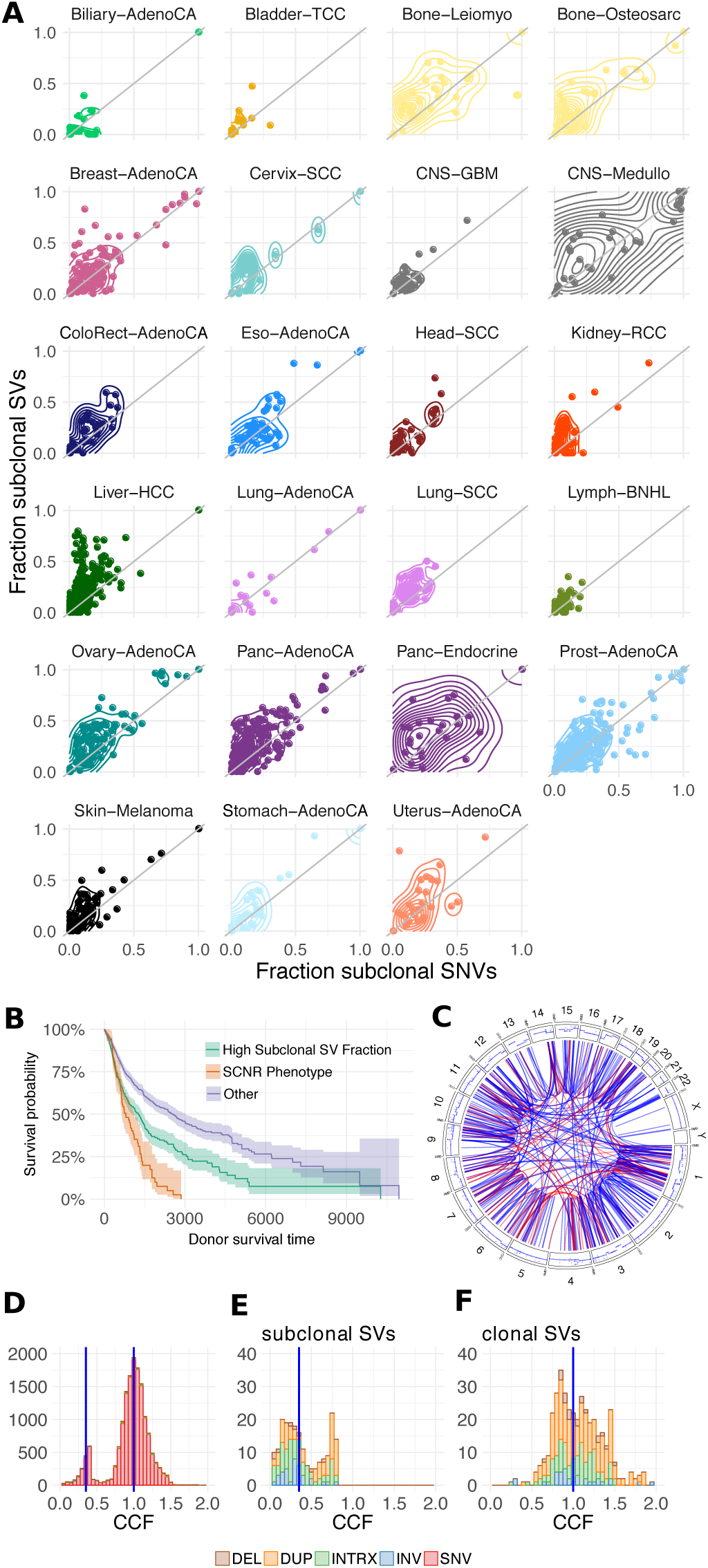
**A)** 2D density plot for PCAWG samples with at least 20 samples per category (n=1,772) (a variant under 0.9 CCF was considered subclonal). **B)** Survival curve comparing SCNR phenotype, SV-enriched and all other PCAWG samples. **C)** Circos plots for example SCNR phenotype tumour (Liver Hepatocellular carcinoma, tumour WGS aliquot 2bff30d5-be79-4686-8164-7a7d9619d3c0). **D)** CCF histogram of sample’s variants. **D)** CCF histogram of SV categories in subclonal cluster. **E)** CCF histogram of clonal SVs.

One unique feature of SVclone is that it determines the clonality of copy-number neutral rearrangements. As such, we applied a test for enrichment of subclonal copy-number neutral rearrangements (inversions and inter-chromosomal translocations) across the PCAWG cohort. A total of 162 samples across 17 cancer types exhibited this novel, subclonal copy-number neutral rearrangement (SCNR) phenotype (e.g. Figure 2c-f), with ovarian (n=31, 14.9% of total ovarian), liver hepatocellular carcinoma (n=27, 6.5% of total liver) and breast cancers (n=24, 10.2% of total breast) overrepresented in this set. To test if SCNR events were the result of a single complex rearrangement event (such as chromothripsis), or were simply a set of unrelated rearrangements, we looked for clustered events, and where possible, attempted to walk the derivative chromosome. SCNR events showed a twofold increase in propensity for being part of a complex event, compared to background (32.5% vs. 18.5%). 50% of these clustered SCNR events were linked by at least one inter-chromosomal translocation, compared to only 13.7% of other samples (see Supplementary Table 4), suggesting these events can span multiple chromosomes. 5.4% of tested SCNR chromosomes could be walked, compared with just 0.6% of background chromosomes. These data suggest that subclonal events present in SCNR samples are likely a result of complex, interrelated rearrangements.

To test for potential clinical relevance of the SCNR phenotype, we compared the overall survival of SCNR cases (n=90), with high subclonal SV fraction cases (n=582), and all remaining cases (n=1070) for which overall survival was recorded. These groups showed significantly different (p<0.001) survival probabilities, with median survival times of 796, 1260 and 2543 days, respectively (Figure 2b). To address the issue of varying background survival rates of different tumour subtypes and different levels of SV enrichment, we stratified on tumour histological subtype and binned tumours on 7 levels of SVs (0, 1-100, 101-200, 201-300, 301-400, 401-500 and 501+). This resulted in a hazard ratio of 1.61 for SCNR cases, significantly higher compared to the baseline cohort (p<0.001). This analysis indicates that considering the clonality of balanced genome rearrangements reveals functionally important and clinically relevant observations. Importantly, considering only the clonality of SNVs and/or SCNAs would have failed to reveal this information.

Despite these successful applications of SVclone, it is important to consider some of its limitations. Our approach infers clusters of SVs with similar cancer-cell fractions but does not infer the phylogenetic history of cellular populations, which is beyond the scope of this contribution. Furthermore, we have simplified our model to consider all breakpoints as independent events despite the fact that in some cases these breakpoints may be part of the same complex SV event. Complex SV types are not identified by SVclone’s classification framework, however, users may specify their own types if known. As more sophisticated methods for classifying complex SV events become available, this could be integrated into the algorithm framework.

In cancers where copy-number neutral rearrangements are common, a significant portion of the clonal landscape has remained, until now, unexplored. Here we have demonstrated a pattern of subclonal variant enrichment that would otherwise go undetected if solely considering the clonality of SCNAs and SNVs. Analysing all variant classes and their respective clonality will ultimately be required to gain a more complete picture of the tumour heterogeneity landscape. We have presented the first integrated software package for modeling the cancer cell fraction of structural variation breakpoints using whole-genome sequencing data and have demonstrated its application in identifying novel patterns of subclonal variation. The software provides a useful additional type of analysis for subclonal quantification, for a more integrated approach to modeling tumour heterogeneity.

## Methods

### Code availability

The SVclone software, user documentation, and example data can be downloaded from https://github.com/mcmero/SVclone. Links to all other source code, including figure generation, can be found in the Supplementary Information.

### Data input

The SVclone algorithm requires at a minimum a list of SV breakpoints and associated tumour BAM file. SV breakpoints can be provided as a VCF or as a tab-delimited file of paired single-nucleotide resolution break-ends. Using an SV caller with directionality of each break-end is recommended. The Socrates^21^ output format is natively supported and allows additional filtering by repeat type and average MAPQ. An associated paired-end, indexed whole-genome sequencing BAM file is required for SV. In the filter step, copy-number information can be added in Battenberg^15^, ASCAT^22^ or PCAWG consensus copy-number formats to aid in correcting VAFs. SNV input is also supported in multiple VCF formats (sanger, mutect, mutect call-stats and PCAWG consensus). Further details of input formats can be found in the repository README file.

### SV classification

We employ a decision-tree based approach based on break-end directionality to classify SV events into six categories: inversions, deletions, tandem duplications, interspersed duplications and intra- and inter-chromosomal translocations. See Supplementary Figures 1 and 2 for the variant types we consider and their associated classification rules, and Supplementary Information 1.3.1 for further details.

### Calculating the number of reads supporting a SV

Each SV considered by SVclone has two genomic locations, denoted here as break-ends. From the BAM file SVclone extracts reads that overlap each break-end and compiles separate counts for: split reads (soft-clipped reads that cross each break-end), spanning reads (read pairs that align either side of the break-end, that do not overlap the breaks) and normal reads (reads that cross or span the break-ends but match the reference). Split reads for both break-end loci (*s_j_*) are summed with the spanning read counts (*c_j_*) to obtain the total supporting read total for the SV (*b_j_*). See Supplementary Figure 3 for how the different read types are counted. To offset aligner-specific behaviour which causes the number of supporting reads to be under-represented, each *b_j_* is subject to a linear adjustment factor that incorporates tumour purity. Adjustment is calculated as 1 + *AF*_supp_ · *π_p_* where *π_p_* is the tumour cell content (tumour purity) and *AF_supp_* is set at 0.12 by default (for data aligned using Bowtie2^23^, if using BWA^24^ we recommend setting this to 0.2). This was inferred from simulations of 100bp short reads with a 300bp mean fragment size. Different adjustment values may be required for different read sizes, insert size distributions and aligners.

### Calculating the number of non-supporting reads

For each SV, normal reads are counted at the break-ends resulting in two normal read count totals (*o_l_*, *o_u_*). Only one of these (*o_j_*) is required for CCF calculation. We choose the normal read count that corresponds to the lowest total copy-number. By selecting the lower copy-number state, the copy-number search space (also known as multiplicity) SVclone must consider when calculating CCF is smaller (see below for details). In cases where one break-end has a clonal background copy-number state, and the other a subclonal state, we preferentially select the side with clonal copy-number. In the case where the SV results in a gain of DNA (interspersed and tandem duplications), the normal read count must be adjusted. We consider the SV classification *κ*_j_ for an SV *j*, where *κ*_j_ ∈ {*DEL*, *DUP*, *INTDUP*, *INV*, *TRX*, *INTRX*}. We define two subsets *κ*_gain_ = {*DUP*, *INTDUP*} where normal reads at the variant population’s break-ends are unaffected at the variant allele, and *κ*_non-gain_ ={*DEL*, *INV*, *TRX*} where the normal reads at the variant population’s break ends are replaced by supporting reads. We compute an adjustment factor,

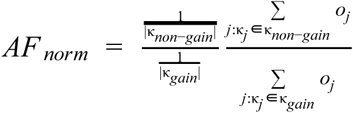

If the tumour contains only SVs in *κ*_gain_, we calculate an approximated *AF_norm_* = 1 − π_⍴_/*n^T^* where π_⍴_ is the tumour content and *n^T^* the tumour ploidy. The normal read counts of all DNA-gain events are then multiplied by this adjustment factor (o_j_ = o_j_ · *AF_norm_* if *κ*_j_ ∈ *κ*_gain_), while events that are not DNA-gains remain unadjusted.

For a more detailed explanation of the read counting, see Supplementary Information 1.4.

### Filtering variants

To obtain a high-confidence set of variants, we apply five filtering criteria: germline variant presence, SV size, minimum depth, minimum support and presence of a valid copy-number state (see Supplementary Information 1.5 for details).

### Assigning copy-number states to SVs

We match SV break loci to the copy-number state of each locus *before* the SV takes place. To do this, we use the SV directionality to determine the flanking copy-number state *outside* the SV, by some offset (100kb by default to allow for segmentation noise). See Supplementary Table 2 for details. Optionally, if the SV does not fall within any CNA boundaries, the nearest bordering CNA state can be assigned (by default no state is matched to the break-end). For further details see Supplementary Information 1.6.

### Clustering

The clustering step of SVclone simultaneously computes SV CCFs and clusters SVs of similar CCF, based on purity, ploidy and copy-number status of the normal, reference and tumour populations. Clustering takes read counts and copy-number states as input and utilises a Bayesian Dirichlet Process mixture model, implemented using Markov-Chain Monte-Carlo (MCMC) sampling (through the PyMC package^25^) in order to approximate posterior distributions for unknown parameters. The software determines the number of clusters dynamically and infers the most likely average CCF per cluster, as well as the multiplicity of each variant (the most likely copy-number states for the tumour’s reference and variant populations, and the most likely allele proportion the variant occurs on). Supplementary Figure S.5 shows the model implementation in graphical model form. Our model relies on the *infinite sites assumption*^26,27^, which states that the same event does not occur twice, independently at the same locus, in descendent populations. We also assume that an SV always occurs on either the major or minor allele, but never on both.

A detailed description of the model can be found in the Supplementary Information 1.7, however, here we highlight the key components. We model the probability of sampling a variant read given variant locus *j* as coming from a binomial distribution with trials *d_j_* (read depth *b_j_* + *o_j_*) and probability *p_j_*,

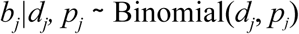

where *b_j_* = *s_j_* + *d_j_*, (*s_j_* is the number of split reads and *d_j_* the number of discordant reads), and *O_j_* the number of normal reads. In order to calculate p_j_ we require the tumour purity estimate π_⍴_ and the subclonal copy-number fraction (φ_*j*_^*bb*^), both of which are specified by the user, along with the unknown parameters cluster CCF mean (*φ_j_*, the proportion of alleles the variant lies on (*μ_j_*) and the total copy-number of the normal (*n_j_^N^*), the reference (*n_j_^R^*) and the variant (*n_j_^V^*) populations:

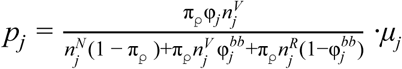

We use MCMC to compute the posterior distribution for *φ_j_* (see below). For, *μ_j_* at each iteration of the MCMC we compute the binomial probability for values of *μ_j_* ranging from 1 to the major allele copy-number over the total variant copy-number (*1*.. *n^V^ _jA_*) / *n^V^ _j_*. and select the value resulting in the *p_j_* with the highest binomial probability.

For detailed information on how *n_j_^N^*, *n_j_^R^*, *n_j_^V^* are computed, see Supplementary Information 1.7.2, Here specify the posterior for φ_j_. In order to avoid fixing the number of subclonal clusters a-priori, we implement a flat Dirichlet Process (DP) with an upper bound *K* on the number of clusters. Hence, φ_j_ is constrained by taking on one of *K* values; φ_j_ ∈ φ_1_..φ_K_. The categorical variable *z_j_* takes on values corresponding to the *K* clusters; *z* ∈ 1, .., *K*. Hence the generative DP model is defined as:

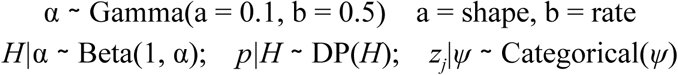

φ’s prior (φ_0_) is defined as the minimum expected prevalence of its associated population, i.e. the minimum φ that can be detected, given that the expected population is present with 1 supporting read on average. The default maximum of φ is defined as 1 (by default).

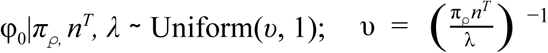

where λ = average depth of coverage.

After the MCMC is complete each variant’s cluster is based on the cluster membership mode across all (post burn-in) iterations. Each cluster’s mean CCF is taken as the average of the post burn-in trace values. Clusters which have no variants assigned are discarded. In addition to the cluster CCF mean, we also provide the post burn-in CCF trace per variant, and a variant CCF calculated from the adjusted VAF (see Supplementary Information 1.7.7). The MCMC is typically run multiple times with a BIC-like fit metric for calculating the model goodness of fit per run (see Supplementary Information 1.7.6).

### Recommended clustering parameters

We recommend using at least 25,000 iterations with 12,500 burn-in, with shape 0.1 and rate 0.5 (for SV data) with 8 runs per sample (default parameters). We recommend lowering the rate parameter to 0.1 when clustering larger numbers of variants (>1000), such as when clustering SNVs. By default, variants are initialised to a single cluster with a clonal CCF. See Supplementary Information 1.7.4 for SVclone’s dynamic initialisation option that pre-clusters data to reduce iterations.

### Post-assignment of variants to clusters

SVclone allows variants filtered out during the filter step, or not present in the clustering step due to subsampling, to be retroactively assigned to the most likely cluster based on their normal and supporting read counts, and their copy-number state. Multiplicity is selected based on the most likely copy-number combination assuming a clonal (φ = 1) state. The most likely cluster from the clustering results is then selected:

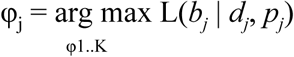

### SV simulation

SVs were simulated with the same tool used to assess the Socrates SV caller^21^, with minor adjustments. SVs size was randomly chosen among the size categories 300-1 kb, 2 kb - 10 kb and 20 kb - 100 kb with equal probability for each category. We simulated chromosome 12 with 100 bp paired-end reads with 300 bp insert size (standard deviation = 20 bp), with an SV at every 100kb interval. Samples containing only deletions, translocations, inversions and duplications were generated at the tumour purity levels of 100%, 80%, 60%, 40% and 20%. The SV events were assumed to always occur in a heterozygous fashion, hence the “true” VAF was always considered to be half of the simulated purity value. To achieve the effect of differing purities, simulated normal reads were mixed with tumour samples with coverage equivalent to (λ · (1 − π_0_)) / 2 and normal read coverage of (λ · π_0_) / 2. We ran simulations at 50x coverage, typical for WGS data by simulating (500*L* / 300) total reads per simulation where *L* is the chromosome length (post rearrangement) and 300 the fragment length. These data were run through SVclone’s annotate (inferring directions), count and filter steps. Adjusted VAFs were used to test the concordance with expected VAF.

### Prostate sample mixing

The metastatic samples bM (A) and gM (B) from Patient 001, Hong et al. were chosen due to their similar coverage (51.5x and 58.9x) and purity (49% and 46%). Previous analysis by Hong et al. showed that these metastases shared a common ancestral clone, had no evidence of subclonality, and contained a number of private SVs and SNVs. Mixing two clonal metastases from the same patient has many advantages over spike-in approaches including: realistic sequencing noise; realistic subclonal mixing of SVs, SCNAs and SNVs; and a natural branching clonal architecture with both clonal and subclonal mutations present. We generated a total of nine samples with subclonal mixes of reads sampled at percentages 10:90; 20:80; 30:70; 40:60; 50:50; 60:40; 70:30; 80:20; and 90:10; for metastasis A and B, respectively. Three clusters are expected to be revealed upon mixing: shared variants present at 100% CCF, one cluster at bM’s mixture frequency and one cluster at gM’s mixture frequency.

These in-silico mixes were created using the subsample and merge functions from SAMtools v1.2^28^. Copy-numbers were obtained from Battenberg^16^ on each merged sample with default parameters. To construct the breakpoint list for input into SVclone’s annotate step, Socrates^21^ was run on the individual bM and gM samples, then run through SVclone’s annotate and count steps (using Socrates’ directions, filtered on simple and satellite repeats using the repeat-masker track (repeatmasker.org) and a minimum average MAPQ of 20). The resulting bM and gM SVs were then merged and filtered against the germline. Copy-numbers were matched using corresponding Battenberg subclonal copy-number output. The merged SV list was used as the set of SV calls for the annotate step for each mix. Clustering was performed across 16 runs with 25,000 iterations, 12,500 burn-in and otherwise default parameters.

The reference and variant alleles were counted at each of the 9811 SNVs across the different mixture proportion BAM files (Mutect variant calls from Hong et al. were used with alleles recounted using Samtool’s mpileup and pileup2base (https://github.com/riverlee/pileup2base). Corresponding Battenberg copy-numbers were matched at each locus, filtering out any variants in regions of subclonal copy-number (PyClone does not support subclonal copy-number handling). For both variant types, we filtered out any clusters that had variant proportions under 5%. Cluster CCFs were derived from the mean φ trace per cluster. We subsampled 5000 variants from the resulting SNV output per mixture and ran these variants through the PyClone algorithm. See Supplementary Figure 4 for full visualisation of the results. See Supplementary Information 1.10 for further details.

### Analysis of ICGC/TCGA pan-cancer samples

We utilised the pan-cancer analysis of whole genomes (PCAWG) October 12^th^ 2016 consensus SNV call set, the v1.6 consensus SVs and the high-confidence 3* annotated copy-number consensus calls (9^th^ of January 2017) using segments with levels a-d as input. For a detailed explanation on how these were generated see the PCAWG-11 heterogeneity manuscript Dentro et al., in preparation. Annotate and count were run using each sample’s associated mini-bam. Consensus purity and ploidy estimates (January 9^th^ 2017) were used. Samples were run for 8 runs, with 25,000 iterations with 12,500 burn-in using a rate parameter of 0.1 (on the DP’s Gamma prior) and otherwise default parameters. SNVs were sub-sampled to 5000 variants, with remaining SNVs post-assigned. SVclone’s custom BIC metric was used to select the best run in each case. Variants falling into clusters with CCF above 0.9 were considered clonal, otherwise subclonal.

We tested the PCAWG samples for the enrichment of balanced rearrangements in subclones (inversions and inter-chromosomal translocation) using a hyper-geometric test, with the alternative hypothesis of
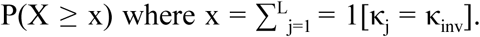
P-values were corrected using false discovery rate (FDR). Survival analysis was undertaken using the *survival* CRAN package (cran.r-project.org/package=survival). Hazard ratios were calculated using the Cox proportional hazards regression model, stratified by tumour histology type. We used a hyper-geometric test to determine whether any ICGC/TCGA contributors were over-represented for each histology type and found no evidence of any significant over-representation (FDR < 0.05). For the clustering of breakpoints criteria, SVs were tested on a per-chromosome basis (inter-chromosomal SVs were removed). Ability to walk each derivative chromosome was tested using criteria for chromothripsis tests A and F^29^. Chromosomes were only tested if they contained at least 4 clonal and 4 subclonal rearrangements per chromosome.

### SVclone software features

SVclone is a modular, flexible and customisable piece of software with over 70 adjustable parameters. The following provides an outline of features:

- SV annotation
  - Classification of event type
  - Inference of SV direction
  - Support for Socrates, GRIDSS and PCAWG consensus SV formats
- SV VAF calculation
  - Normal supporting spanning and split read counts
  - Break-end matching to background copy-number (given SCNA states in Battenberg, ASCAT or PCAWG consensus formats)
  - Selection of break-end side for normal read count
- Filtering and adjustment
  - Filtering criteria include germline presence, SV size, read count (depth, minimum split and spanning reads), overlap of bed regions to exclude and background copy-number state (optionally filter subclonal states or non copy-number neutral)
  - Correction of DNA-gain normal reads via adjustment factor
  - Correction of supporting reads by scalable factor
  - Support for SNVs in mutect, mutect callstats, sanger and PCAWG consensus formats
- Clustering
  - Support for SVs and SNVs (option to cocluster, or cluster separately)
  - SNVs may be subsampled
  - Automatically infer a fixed alpha based on number of variants
  - Inference of cluster number
  - Cluster number initialisation based on power to detect (can also be manually set)
  - Clusters can be merged based on confidence interval range
  - Increase multiplicity search space to consider more copy-number combinations
  - Model for estimating most likely SCNA states for variant and reference populations
    ◼ Option to weight clonal copy-number states more strongly
  - Correction for CCF traces due to label switching
  - Calculation of fit metric for selecting best run
- Post-assignment
  - Filter out small clusters
  - Variants can be (re)assigned to their most likely cluster. These may be:
    ◼ variants from filtered out clusters
    ◼ variants used in the cluster step
    ◼ variants not used in the cluster step
- Post run metrics (See Supplementary Figure 5 for summary of output plots)
  - Circos plots (including SVs, copy-number and SNV density)
  - Overview plot of clustering results and model fit across runs
  - Histogram plots per run (VAFs and CCFs by cluster and variant type)
  - Coclustering matrix output

